# The contrasted impacts of grasshoppers on soil microbial activities in function of primary production and herbivore diet

**DOI:** 10.1101/2022.07.04.497718

**Authors:** Sébastien Ibanez, Arnaud Foulquier, Charles Brun, Marie-Pascale Colace, Gabin Piton, Lionel Bernard, Christiane Gallet, Jean-Christophe Clément

**Affiliations:** Univ. Grenoble Alpes, Univ. Savoie Mont Blanc, CNRS UMR 5553, LECA, Chambéry, France; University of Applied Sciences and Arts Western Switzerland – Land, Nature, Environment Institute, Hepia Geneva, Route de Presinge 150, CH-1254, Jussy, Switzerland; Univ. Savoie Mont Blanc, INRAE, CARRTEL, Thonon-Les-Bains, France

**Keywords:** nutrient cycling, primary productivity, grasshoppers, grasslands, ecosystem functioning, plant-soil feedbacks

## Abstract

Herbivory can have contrasted impacts on soil microbes and nutrient cycling, which has stimulated the development of conceptual frameworks exploring the links between below- and aboveground processes. The “productivity model” predicts that herbivores stimulate microbial activities and accelerate nutrient mineralization in productive ecosystems, while they have an opposite effect in less productive ecosystems. In parallel, the “diet model” predicts that herbivores feeding on conservative plants accelerate nutrient cycling while those feeding on exploitative plants decelerate nutrient cycling, due to changes in litter inputs. Since these two frameworks can lead to conflicting predictions in some cases, experimental evidence combining herbivore diet and plant productivity is required.

During two consecutive years, we conducted an experiment controlling the presence of three grasshopper species consuming either grasses, forbs or both in twelve natural and managed alpine grasslands with contrasted productivities. In order to assess the effects of herbivory on soil microbes, we measured their extracellular enzymatic activities, biomass and potential nitrogen mineralization (PNM). Soil and vegetation were also characterized to test how much they modulated the effects of herbivory on microbes.

Contrary to the predictions of the diet model, the effects of herbivory on microbial characteristics did not depend on the herbivores diet, but were influenced by primary production, though in a way that differed from the productivity model. The most productive sites were constituted by exploitative plant species which depleted N resources in the soil, and by microbes producing relatively few extracellular enzymes, leading to a lower PNM. Herbivory increased microbial biomass and decreased the production of extracellular enzymes in those sites, possibly through the stimulation of root exudates produced by exploitative species. The least productive sites were characterized by conservative plants, high soil C content, and by microbes having a resource acquisition strategy (more extracellular enzymes, higher PNM). Herbivory decreased microbial biomass and increased the production of extracellular enzymes in those sites. This pattern can be explained by the loss of carbon associated with insect respiration, which increases the resource requirements of microbes and by a lower production of root exudates by conservative species. Therefore, the effects of two years of herbivory on soil microbes were at odds with the productivity model, which focuses instead on longer term effects corresponding to herbivory-induced changes in plant species composition. This highlights the multidimensional feature of the impacts of herbivory on ecosystem functioning, both in space and time.

## Introduction

During the last decades the influence of herbivory on terrestrial ecosystem functioning has been highlighted through its effects on matter and energy fluxes linking above and belowground communities (Parker et al. 2017; Kristensen et al. 2020; Sandén et al. 2020). Plant-herbivore interactions influence the quantity and quality of organic matter inputs to soil detrital food webs with important implications on the rate of microbial processes which regulate organic matter decomposition and nutrient recycling and ultimately control the maintenance of soil fertility and carbon sequestration (Bardgett and Wardle 2010). Previous studies have already reported a diversity of impacts of herbivory on soil carbon and nutrient cycling (Hunter 2001; Bakker et al. 2004), including positive (e.g. Frank et al. 2000; Belovsky and Slade 2000), negative (e.g. Ritchie et al. 1998) or no detectable effects (Singer and Schoenecker 2003). An overarching conceptual model of these contrasting effects is needed to understand which ecological variables control the direction and magnitude of the effects of herbivory on soil microbial communities and related ecosystem processes.

Among the different frameworks that have been proposed, one focuses on the contrasting effects of ecosystem productivity (Bardgett and Wardle 2003, 2010; Wardle et al. 2004), and one on the mitigation by herbivore diet (Ritchie et al. 1998; Belovsky and Slade 2000; Hunter 2001; Tuomi et al. 2019). The “productivity model” predicts that in productive ecosystems herbivores consume a high percentage of the net primary production, rapidly returning organic matter to the soil as easily decomposable fecal material enriched in nutrients (“fast cycle”, McNaughton et al. 1988). Herbivores also promote compensatory plant growth (McNaughton 1983) or nutrient reallocation in leaf tissues of exploitative plant species (Potthast et al. 2021), which have high leaf nutrient content (low C/N) and grow faster. Herbivores also slow down the establishment of conservative plant species (high C/N, slow growing) which produce more recalcitrant litter (Reich 2014). The combination of these positive effects on the quality of detrital resources leads to an acceleration of nutrient cycling, which further induces a positive feedback loop. Instead, in less productive ecosystems herbivores consume a smaller proportion of the net primary production, favoring the accumulation of recalcitrant plant litter (“slow cycle”, McNaughton et al. 1988). This comes along with the promotion of conservative plant species producing even more recalcitrant litter therefore promoting a slower nutrient cycling.

In contrast, the “diet model” focuses on another aspect of herbivore-plant-soil interactions and distinguishes two types of herbivores, those consuming exploitative plants, and those eating conservative plants. Indeed, although most vertebrate herbivores either feed non-selectively or prefer high-quality plants (Hofmann 1989; Clauss et al. 2003), many insects prefer tougher plants (Ibanez et al. 2013) and perform better on seemingly low-quality diets (Cease et al. 2012; Talal et al. 2020). Herbivores feeding on exploitative plants favor the growth and survival of conservative plants and therefore the accumulation of more recalcitrant litter. This slows down organic matter cycling, as predicted by the productivity model according to which herbivores preferentially feed on higher quality plants in the least productive ecosystems (Bardgett and Wardle 2010). Instead, herbivores feeding on conservative plants transfer low quality litter into the fast cycle, which accelerates decomposition and nutrient cycling, and promotes exploitative plant species (Belovsky and Slade 2000). During the last 20 years, several experiments controlling for herbivory either by ungulates or insects provided evidence in favor of the diet model. When herbivores fed on the most conservative plants, they increased primary production and/or nutrient availability (McNaughton et al. 1997; Belovsky and Slade 2000, 2018; Garibaldi et al. 2007; Schmitz 2008; Nitschke et al. 2014). In contrast, when herbivores fed on the most exploitative plants they had an opposite effect (Pastor et al. 1993; Ritchie et al. 1998; van Wijnen et al. 1999; Harrison and Bardgett 2004; Schmitz 2008; Belovsky and Slade 2018). The two experiments showing such combination of both effects used polyphagous grasshopper species for which the diet changes either according to the type of predators present (Schmitz 2008) or depending on intraspecific variation of leaf water content (Belovsky and Slade 2018). However, these experiments as well as others (e.g. Deraison et al. 2015) manipulated either the presence or the diets of herbivores in a single type of ecosystem, which hampers the articulation of the diet model with the productivity model.

Although the productivity and diet models focus on different aspects (ecosystem productivity and herbivore diet), they can be combined. In some cases, the predictions of both models are aligned (Table 1). However, in less productive ecosystems inhabited by herbivores selecting low quality plants, nutrient cycling may be affected in both directions, depending on the relative strength of the various processes at play in the productivity and diet models. Similarly, herbivores feeding on high quality plants in fertile ecosystems may have contrasting effects on matter turn-over (Table 1). For instance, under fertile conditions herbivores feeding on high quality plants can favor their dominance (Buckland and Grime 2000), presumably because high resource availability allows compensatory growth for these plant species. This may in turn promote nutrient cycling and soil fertility (Bardgett and Wardle 2010). However, if high-quality plants invest in defensive secondary compounds instead of displaying compensatory growth, and if the herbivores can cope with them, slow-growing species may become dominant and nutrient cycling may slow down.

**Table 1.** Expected effects of herbivory on matter cycling (either accelerating or decelerating), according to the productivity and diet model. In each of the four combinations, the prediction of the diet model is reported on the top (in orange), and that of the productivity model on the bottom (in green). In two cases out of four, the predictions are opposite.

|  |  | Diet model |  |
| --- | --- | --- | --- |
|  |  | High quality diet | Low quality diet |
| Productivity<br>model | Productive ecosystems (high plant biomass,<br>low soil organic matter) | decelerate | accelerate |
|  |  | accelerate | accelerate |
|  | Less productive ecosystems (low plant<br>biomass, high soil organic matter) | decelerate | accelerate |
|  |  | decelerate | decelerate |

Furthermore, while the productivity and diet models explicitly consider plant and herbivore resource acquisition strategies, they do not consider the resource acquisition traits of soil microbes. Yet, microbial communities are key regulators of nutrient recycling in the plant-soil system, and recent developments suggest that including microbial communities in a multitrophic functional trait framework could strengthen our mechanistic understanding of ecosystem functioning (Malik et al. 2020; Piton et al. 2020b). Resource acquisition strategies of microbial communities can be partly inferred from the characterization of enzymatic activities involved in C, N and P acquisition through the extracellular depolymerization of organic compounds (Piton et al. 2020b, a). A higher investment in resource acquisition traits (i.e. increased extracellular enzyme activities) is expected along a gradient of decreasing soil resource availability, paralleling a shift in plant resource acquisition strategy along the exploitative-conservative continuum (Piton et al. 2020b). Increased quantity (i.e. soil organic matter, N and P amounts) and quality (i.e. lower soil C/N, higher nitrate, ammonium contents) of available resources from directly assimilable fecal material enriched in nutrients (e.g. Fielding et al. 2013) or increased labile carbon inputs from root exudates (Hamilton and Frank 2001; Paterson et al. 2003; Hamilton et al. 2008) are expected to promote microbial communities (Wardle et al. 2004; Grigulis et al. 2013) with a high yield strategy (Y strategy, Malik et al. 2020) characterized by a low investment in extracellular enzyme production (Piton et al. 2020b). On the contrary, a reduction of available resource quality should promote microbial communities with a resource acquisition strategy characterized by a high investment in extracellular enzymes targeting complex polymeric organic matter (A strategy, Malik et al. 2020). Higher nitrogen availability in the form of labile compounds (amino acids, NH_4^+^_ or NO_3^-^_) resulting from herbivory should reduce the microbial demand for this element, resulting in a lower investment in N-acquiring enzymes. Yet, increased availability of labile C from root exudates following herbivory might also increase nutrient limitation of microbial communities, especially if herbivory stimulates plant productivity and associated nutrient acquisition. This would result in higher nutrient immobilization, which is more likely under less fertile conditions that offer low resource availability for microbes. Moreover, the productivity model focuses on the contrasting effects of herbivory along a gradient of plant productivity, but soil nutrient cycling is ultimately controlled by microbial communities, whose responses to herbivory may rather depend on a gradient of soil resources, on which they rely more directly than on plant productivity. Both gradients are often correlated, productive ecosystems being associated with low carbon content (Wardle et al. 2004; Grigulis et al. 2013), but decoupling can also be observed (Piton et al. 2020b).

We therefore identified two important gaps in previous works regarding the effects of herbivory on ecosystem functioning: first, the two existing frameworks should be combined (Table 1), and second, the functional characteristics of soil microbial communities should explicitly be included within the frameworks. This calls for experimental evidence controlling for herbivore diet in ecosystems with contrasting plant productivity and soil resource levels, in order to test the effect of herbivory on microbial communities. In the present contribution, we bridge these gaps with a semi-controlled experiment using insect-proof cages, where four types of grasslands (managed or natural, dominated by grasses or by forbs) in 12 montane sites were subjected to herbivory by three grasshopper species which either preferred grasses, forbs or a mixture of the two. Since within each site grasses had a higher C/N ratio than forbs (Ibanez et al. 2021), grass-feeders were expected to accelerate matter cycling, while an opposite effect was predicted for forb-feeders. Following two years of herbivory, activities of extracellular soil enzymes specialized in carbon, nitrogen and phosphorus acquisition were measured, as well as nitrogen mineralization potential and microbial biomass. We hypothesized that the effect of herbivory on these soil microbial characteristics depends on herbivore diet and plant productivity. In cases where the two frameworks make opposite predictions (Table 1), the experiment was designed to compare the relative importance of diet and productivity effects.

## Material & methods

### Study sites and experimental design

The experiment was conducted in the French Pre-Alps, in Northern Vercors on the calcareous plateau of Autrans-Méaudre (45°10’N, 5°32’E). Four grassland types were chosen on the base of their botanical composition: extensively managed and fertilized grasslands dominated by the grass *Festuca rubra*, intensively managed and fertilized grasslands dominated by the grass *Lolium perenne*, natural grasslands dominated by forbs characterized by *Heracleum sphondylium* and *Chaerophyllum hirsutum* and natural warm grasslands mainly formed by the grass *Bromopsis erecta*. The 6 managed grasslands are disseminated on the Autrans plateau, and the 6 natural grasslands form two distinct clusters containing either 2 forbs-dominated communities and 1 warm grassland, or 1 forbs-dominated community and 2 warm grasslands (Sup. Fig. 1, site coordinates in Sup. Tab. 1). Since the two clusters contain communities having contrasting plant productivity, botanical composition and soil characteristics, spatial autocorrelation is unlikely to affect the results. All grasslands were composed of a mixture of forbs and grasses, although the dominant functional group depended on the grassland type. Their productivity ranged from 156 to 840 g dw yr-1 m-2, managed grasslands being more productive than natural grasslands (Sup. Fig. 2). Three replicates per grassland type were randomly chosen in the study area, for a total of twelve sites ranging from 1000 to 1300 m.a.s.l. In each site, five 1m^2^ plots (60 plots in total) were selected randomly within a 5m-radius and assigned to one of the five following grasshopper herbivory treatments: *Miramella alpina*, *Pseudochorthippus parallelus*, *Stauroderus scalaris*, an equal mixture of the three species, no grasshoppers (control). If single species treatments have contrasted effects on soil microbes, the mixture treatment should discriminate between additive and non-additive effects. The grasshopper species were chosen according to their diet, as *M. alpina* prefers to feed on forbs, *P. parallelus* on grasses and *S. scalaris* mainly on grasses but also on forbs (Ibanez et al. 2013). Plots were covered by a 1m^3^ metallic cage covered by insect-proof mesh (PE 22.30, 920 × 920 μm; DIATEX, Saint Genis Laval, France), into which the adult grasshoppers were introduced. The number of grasshoppers per cage was adjusted to fit 1-g of grasshopper for 300-g of plant aboveground biomass (dry mass). Preliminary field observations revealed that this ratio corresponds to the maximal natural insect densities locally observed in the study area. This standardization is necessary to avoid confounding correlations between herbivory load and herbivore diet, or between herbivory load and plant productivity. For the standardization, the dry mass of each species and each sex was also considered, after preliminary measurements. Grasshoppers were present inside the cages during 65 days from the 2^nd^ week of July until the 3^rd^ week of September, during two consecutive years 2016 and 2017. During these periods, the number of grasshoppers per cage was checked every two weeks, including the control cages from which a few grasshoppers were occasionally removed. Several ecosystem characteristics were measured, as detailed in the following sections. In short, soil abiotic and microbial characteristics as well as plant biomass were measured at the end of the experiment, in mid-September 2017. Herbivory marks on standing biomass was visually estimated in 2016, while plant specific abundance was estimated before the experiment in 2016 and after one year in 2017, in order to calculate the effect of herbivory.

### Soil characteristics

In mid-September 2017, four 10-g soil samples were collected in each plot at 5-cm depth, and then bulked. The samples were sieved at 5-mm and stored at 4°C before processing within 48h. Subsamples of 5-g of soil were dried at 70°C for 1 week to determine soil moisture, followed by 4h at 550°C to determine soil organic matter (SOM) content. Soil subsamples were air dried and ground to powder to measure total C and N contents using a FlashEA 1112 elemental analyzer (Fisher Scientific Inc., Waltham, MA, USA), and to determine soil pH in a 1:2.5 (soil: distilled water) solution. Solution of 0.5M K_2_SO_4_ was used to extract soil nitrate (NO_3^-^_), ammonium (NH_4^+^_), total dissolved nitrogen (TDN) (Jones and Willett 2006), and phosphate (PO_4^-^_) on 10-g of fresh soil. N and P concentrations were measured on an automated photometric analyzer using standard colorimetric methods (Gallery Plus: Thermo Fisher Scientific, Waltham, Massachusetts, USA). Dissolved organic nitrogen (DON) was calculated as the difference between TDN and the mineral N (NO_3^-^_ + NH_4^+^_).

### Microbial biomass, activities and traits

The same soil samples were used for the analysis of microbial characteristics. Microbial biomass nitrogen (MBN) content was based on the difference of soil N content before and after chloroform-fumigation extraction of 10 g of fresh soil (Vance et al. 1987). We then calculated MBN in μgN/g using a correction factor of 0.45 (Jenkinson et al. 2004). Potential nitrogen mineralization rates (PNM) were estimated after incubation under anaerobic conditions of 10 g of fresh soil for 7 days at 40°C in the dark (Wienhold 2007). Mineralized organic N was accumulated as NH_4^+^_ during this incubation and PNM rates were calculated based on the difference between NH_4^+^_ content before and after incubation and expressed as μgN/g dry soil/day. Microbial resource acquisition strategies were characterized using different microbial community-weighted mean traits (Piton et al. 2020b). We measured the potential activity of seven extracellular enzymes contributing to the degradation of C-rich substrates (α-glucosidase, β-1,4-glucosidase, β-d-cellobiosidase and β-xylosidase), N-rich substrates (β-1,4-N acetylglucosaminidase and leucine aminopeptidase), and P-rich substrates (phosphatase) using standardized fluorimetric techniques (Bell et al. 2013). We homogenized 2.75 g of fresh soil (1 minute in a Waring blender) in 200 ml of sodium acetate buffer solution that was adjusted at the mean soil pH observed in the present study (i.e. 6.2). The resulting soil slurries were added in duplicates to 96-deep-well microplates. We then added a substrate solution for each enzyme. Duplicated standard curves (0–100-μM concentration) were prepared for each soil sample by mixing 800 ml of soil slurry with 200 ml of 4-methylumbelliferone (MUB) or 7-amino-4-methylcoumarin (MUC) in 96-deep-well microplates. Microplates were incubated during 3hrs (dark, 175 rpm, 20°C), and centrifuged at 2,900 g for 3 min. Soil slurries (250 μl) were then transferred into black Greiner flat-bottomed microplate and scanned on a Varioskan Flash (Thermo Scientific) reader using excitation at 365 nm and emission at 450 nm (Bell et al. 2013). For each soil sample, the four enzyme activities degrading C-rich substrates, the two enzymes activities degrading N-rich substrates and all the seven enzymes were summed to obtain extracellular enzyme activity for C-rich substrates (EEC), N-rich substrates (EEN) respectively. Phosphatase activity was used to represent extracellular enzyme activity for P-rich substrates (EEP). EEC, EEN and EEP were calculated per gram of dry soil per hour (global activities, nmol activity g-1 dry soil h-1). Microbial resource limitation and associated trade-offs between C and N acquisition was assessed through the calculation of the ecoenzymatic ratio EEC:EEN (Sinsabaugh et al. 2009).

### Vegetation surveys and plant biomass

In order to quantify the diet of each grasshopper species, a visual estimation of herbivory intensity was conducted in August 2016. Ten individuals belonging to each of the 6 most dominant plant species in each site were inspected for herbivory marks by grasshoppers and the percentage of leaf area eaten was visually estimated.

A botanical survey was conducted at the beginning of the experiment (June 2016) and after one year of controlled herbivory (June 2017). The point quadrat method was used (Levy and Madden 1933; Vittoz and Guisan 2007; Lavorel et al. 2008), with 50 regularly spaced points in each 1m^2^ cage. This method allows to determine the relative abundance of the most common species in each plot, but does not constitute a complete inventory of the specific richness. To assess the effect of herbivory, we computed the proportion of forbs in 2017 minus the proportion of forbs in 2016 in each cage. At the end of the experiment in September 2017 the aboveground biomass was harvested in each plot, dried 48h at 40°C, sorted into forbs and grasses and weighed. In order to get a community-level measure of leaf C/N ratio, a representative sample of the harvested biomass was ground to powder, homogenized, and 5 mg of the leaf powder were then analyzed for carbon and nitrogen concentration using a CHN analyser (FlashEA 1112, ThermoFisher Scientific, MA, USA). These measures also provided the amount of nitrogen in aboveground plant biomass per unit area (gN/m^2^, N_B_).

### Statistical analysis

Linear mixed models in which the random effects corresponded to the 12 sites (with 5 pseudo-replicas per site) were used to predict the logit of the total percentage of leaf biomass eaten in function of grassland type, herbivore identity, plant functional group (forbs *vs* grasses), and the interaction between these three factors.

Coinertia analysis (Dray et al. 2003) was used to quantify the coefficient of correlation (RV coefficient, ranging between 0 and 1) between vegetation characteristics (total aboveground biomass, forbs biomass, N_B_, community-level plant C/N ratio, plant Shannon index), soil abiotic characteristics (water content, pH, SOM, TDN, phosphorus content, C/N ratio) and soil microbial characteristics (EEC, EEN, EEP, EEC:ENN ratio, PNM and microbial biomass). Redundancy analysis (RDA, Borcard et al. 1992) was used to quantify the amount of variation of either microbial, soil abiotic and vegetation characteristics explained by either grassland type or site.

In order to quantify the effect of herbivory on the six microbial characteristics, the standardized response to herbivory was calculated as the difference between grasshopper treatments and the control treatment (no grasshoppers) of the same site, divided by the standard deviation of the corresponding site. Coinertia analysis was used to test if the standardized responses depended on soil abiotic and/or on vegetation characteristics.

To test if these effects of herbivory depended on grassland characteristics (either SOM or plant productivity N_B_), and on herbivore identity, the standardized responses of microbial characteristics were then used as a response variable in separate linear mixed models including sites as random effects. Since the warm grassland #1 had a particularly high SOM, the same models as above were performed excluding this site to check if it had a disproportionate effect on the results.

In all mixed models, the denominator degrees of freedom were calculated using Satterthwaite’s approximation (Satterthwaite 1946). Type III sums of squares were used for models including interactions between factors. For each model, the normality and homoscedasticity hypothesis were visually checked. All statistical analyses were performed with the R software (R Core Team 2019), using the packages lmerTest (Kuznetsova et al. 2017) and ade4 (Dray and Dufour 2007).

## Results

### Leaf quality and consumption

The percentage of leaf area eaten depended to a large extent on the interaction between plant functional group and herbivore identity (p<0.001), indicating that the selected grasshoppers had contrasted diets as expected. *M. alpina* preferred forbs over grasses (7% of leaf area consumed *vs* 3.2%), *S. scalaris* preferred grasses over forbs (7.2% *vs* 3%), while *P. parallelus* ate almost exclusively grasses (11.3% *vs* 0.9%, Sup. Fig. 3A). When these three species were combined, grasses were more impacted than forbs (7.4% *vs* 4.8%). The overall effect of grassland type on the leaf area eaten was not significant (p=0.50), which reflects the fact that the number of introduced grasshoppers in each cage depended on the estimated plant standing biomass (1-g of grasshopper for 300-g of plant aboveground biomass). However, there was a significant interaction between grassland type and plant functional group (p<0.001), because in forb dominated communities grass species were much more heavily impacted than forb species (15.3% *vs* 2.7%), whereas in the other grassland types both functional groups were consumed in similar proportions (Sup. Fig. 3B). The p-value of the three-way interaction between plant functional group, grassland type and grasshopper identity equaled 0.58, which indicates that the diet of each grasshopper species did not depend on grassland type. The elementary analysis of fresh plant material showed that the leaf C/N ratio was higher for grasses than for forbs, whatever the grassland type (p<0.001 in all cases). However, the magnitude of the difference depended on grassland type (interaction between plant functional group and grassland type, p<0.001). The largest difference was in warm grasslands (forbs: 25, grasses: 38) and the smallest in communities dominated by forbs (forbs: 22, grasses: 27, Sup. Fig. 4). The botanical survey conducted after one year of herbivory revealed that the proportion of forbs did not vary in most treatments (p>0.4), except in the cages containing the forb-feeding *M. alpina* where forbs declined by 7.2% (Sup. Fig. 5, p=0.013).

### Relations between vegetation, soil microbial and abiotic characteristics

The coinertia analysis between six microbial soil characteristics (nitrogen mineralization potential, EEN, EEC, EEP, microbial biomass, EEC/EEN ratio) and six abiotic soil characteristics (water content, pH, SOM, TDN, phosphorus content, C/N ratio) had a RV coefficient of 0.59 (p<0.001). One axis of covariation between the two types of variables corresponds to an anticorrelation between soil pH and soil phosphorus content on the one hand, and EEP on the other hand, as well as EEC/EEN ratio to a lesser extent (Figure 1A). Therefore, in acidic soils having low phosphorus content, microbes invested more in phosphorus acquisition enzymes and slightly more on carbon than on nitrogen acquisition enzymes. Another axis of covariation was between the abiotic characteristics TDN, SOM and water content, and the microbial characteristics EEC, EEN and PNM. The RDAs indicated that grassland types were heterogeneous with respect to soil characteristics, grassland type explaining 28% of the variation for the six soil abiotic characteristics but only 9% for the six microbial characteristics. In contrast, the soil characteristics were homogeneous at the site level (Sup. Fig. 6).

**Figure 1.**
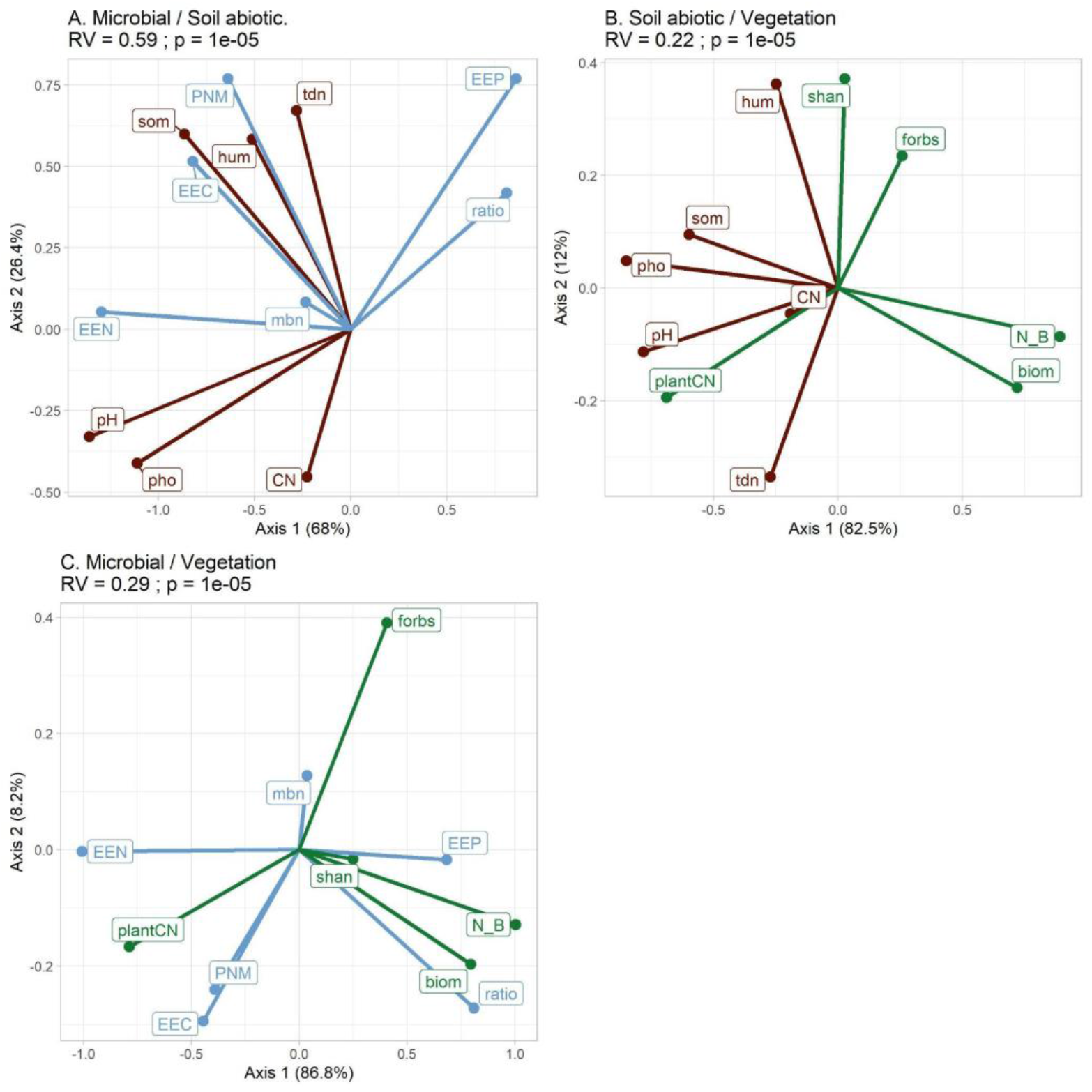
Coinertia of (A) soil microbial and abiotic characteristics, (B) soil abiotic and vegetation characteristics and (C) microbial and vegetation characteristics of the 60 cages. In brown, soil abiotic characteristics (tdn=total dissolved nitrogen, hum=soil water content, som=soil organic matter, CN=soil C/N ratio, pho=soil phosphorus content). In blue, soil microbial characteristics (PNM=potential nitrogen mineralization, mbn=microbial biomass, EEN, EEC, EEP= extracellular enzyme activities related to either nitrogen, carbon or phosphorus, ratio=EEC/EEN). In green, vegetation characteristics (plantCN=community-level plant C/N ratio, biom=total aboveground plant biomass, forbs=forbs biomass (g), N_B=nitrogen content in total aboveground plant biomass (N_B_), shan=Shannon diversity index of plants). The % indicate the projected inertia.

Turning to vegetation, the coinertia analysis between the six abiotic soil characteristics and the five vegetation characteristics (total aboveground biomass, forbs biomass, N_B_, community-level plant C/N ratio, plant Shannon index) had a RV coefficient of 0.22 (p<0.001). Ecosystem productivity was negatively associated with SOM, pH and soil phosphorus content, and was not linked to TDN (Figure 1B). The RV coefficient of the coinertia between five vegetation characteristics and the six microbial characteristics was equal to 0.29 (p<0.001). This much lower RV coefficient than that between soil microbial and soil abiotic characteristics (0.59) suggests that the microbial soil characteristics were more related to soil abiotic than to vegetation parameters. The EEC/EEN ratio covaried with plant productivity (aboveground biomass and N_B_) while EEC, EEN and PNM were associated with the leaf C/N ratio and the proportion of grasses (Figure 1C). The RDA indicated that grassland types were homogeneous with respect to vegetation characteristics with 53% of variation explained by grassland type (Sup. Fig. 6). Indeed, warm grasslands were characterized by high leaf C/N ratio, forb-dominated grassland by a high proportion of forbs and a high Shannon diversity, intensive grasslands by a high biomass and nitrogen content, while extensive grasslands had intermediate vegetation characteristics. Means ± sd of each vegetation, soil microbial and abiotic characteristics are given for each of the 12 sites in Sup. Tab. 1.

### Effect of herbivory on microbial characteristics

Another coinertia analysis, this time involving the standardized responses of microbial communities to herbivory, revealed that they were related to soil abiotic characteristics (RV=0.22, p<0.001). The coinertia analysis (Figure 2) and the linear mixed models (Figure 3) both indicated that the responses of PNM and EEN to herbivory increased with SOM, as well as EEC and EEP although to a lesser extent (see Figure 3 for the p-values of the linear mixed models). More specifically, below 10% of SOM herbivory decreased PNM and soil enzymatic activities while above 15% of SOM herbivory had a positive effect on these variables (Figure 3). The response of microbial biomass to herbivory did not depend on SOM (p=0.75) but varied according to TDN. In sites with TDN below 30 μgN/g, herbivory increased microbial biomass, while in sites with TDN above 50 μgN/g herbivory decreased microbial biomass (Figure 3E, p=0.014). Finally, the response of EEP to herbivory was positively related to pH (p=0.007). In contrast, the effect of herbivory on microbial characteristics did not depend on vegetation traits (RV=0.09, p=0.17). In particular, neither plant biomass nor productivity (N_B_) were related to changes in the standardized responses of microbial characteristics (details not shown, p>0.05). In all cases, the effects of herbivory on microbial characteristics depended neither on herbivore diet nor on the interaction between herbivore diet and ecosystem characteristics such as SOM, TDN or productivity (details not shown, p>0.05).

**Figure 2.**
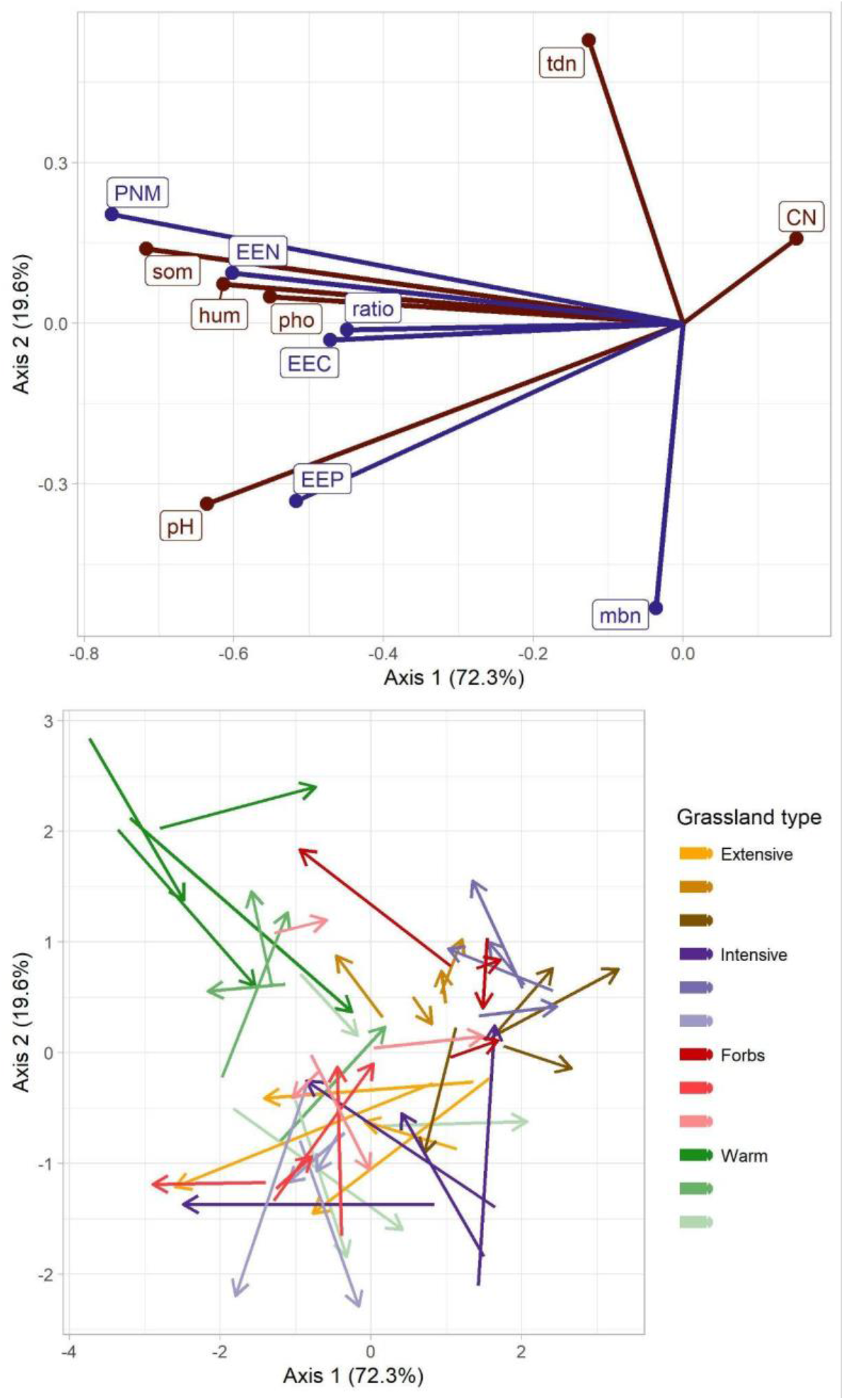
Coinertia of soil abiotic characteristics and the standardized response to herbivory of microbial characteristics. (A) Canonical weights of soil abiotic characteristics (in brown) and standardized response to herbivory of microbial characteristics (in purple). (B) The 48 arrows (60 cages - 12 controls) correspond to all the plots containing grasshoppers. The soil abiotic characteristics are at the beginning of the arrows, the standardized response to herbivory of microbial characteristics are at their end. tdn=total dissolved nitrogen, hum=soil water content, som=soil organic matter, CN=soil C/N ratio, pho=soil phosphorus content, PNM=potential nitrogen mineralization, mbn=microbial biomass, EEN, EEC, EEP= extracellular enzyme activities related to either nitrogen, carbon or phosphorus, ratio=EEC/EEN. The % indicate the projected inertia.

**Figure 3.**
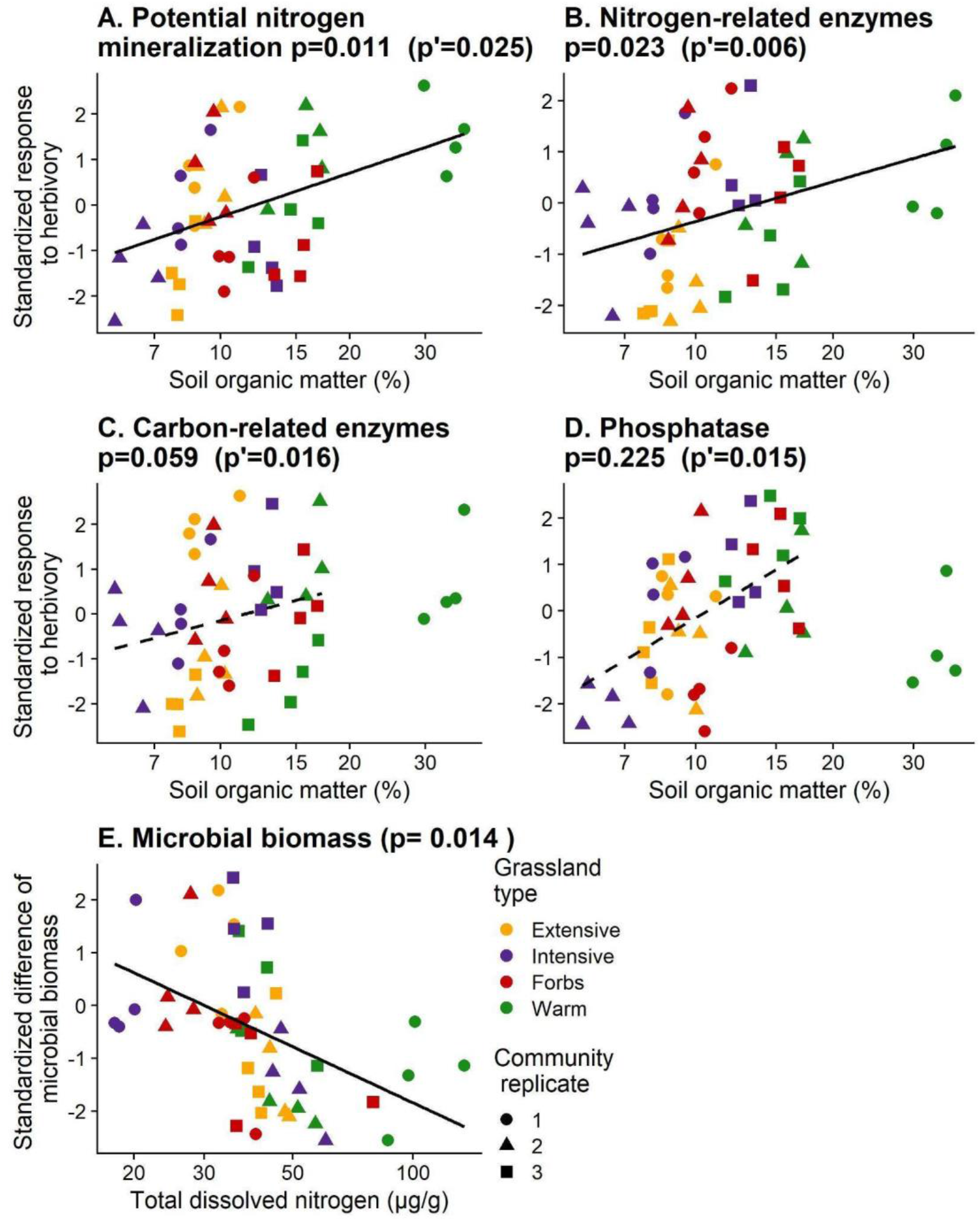
Relationship between soil organic matter and the standardized response to herbivory, calculated as the difference between each grasshopper treatment and the control treatment (no grasshoppers) of the same site, divided by the standard deviation of this site (n=48, 60 cages - 12 controls). p is the p-value of the linear mixed model including all sites (continuous regression line), while p’ excludes the warm grassland #1 having SOM values >30 (dashed regression line).

## Discussion

### Covariations between ecosystem properties

We first explore the covariations between vegetation, soil biotic and soil abiotic characteristics, in order to better identify the ecological factors which differentiate the studied grasslands. In the next sections, we discuss how these differences condition various types of responses to herbivory. The comparison of the 12 study sites indicated that, in line with the productivity model, soil C content measured by SOM was low in sites having the higher plant biomass (Figure 1B). However, there was a positive relationship between SOM and TDN (Figure 1A), and no relationship between TDN and plant biomass (Figure 1B). The decoupling between TDN and plant biomass might be explained by management, since extensive and intensive grasslands are fertilized with manure each year; intensive grasslands being the most fertilized and productive ones in the study area (Piton et al. 2020b & Sup. Fig. 2). Nutrients available after fertilization are likely to be rapidly assimilated by the fast growing and productive plant species, resulting in both high productivity and low measured nutrient availability. After plant growth, mowing exports nutrients away from these grasslands, while fertilization reintroduces the lacking nutrients. Therefore, in managed grasslands a high proportion of annual primary production is consumed and indirectly returned to the soil through subsequent fertilization, as are natural fertile ecosystems in the productivity model (Bardgett and Wardle 2010).

Turning to microbes, those living in soils with high SOM and TDN accelerated the nutrient cycle (high PNM, EEC and EEN, Figure 1A). However, microbial biomass did not increase with SOM, which implies that high SOM comes along with higher mass-specific extracellular activities for carbon uptake and higher mass-specific nitrogen mineralization, which corresponds to a resource acquisition (A) strategy at the expense of growth yield or carbon use efficiency (sensu Malik et al. 2020). Microbial biomass did not covary with any soil parameters (Figure 1 A & C), as in Farrell et al. (2011, see their Table 1) and Chu & Grogan (2010, see their Tables 1 & 2). This may reflect the fact that microbial biomass results from the complex interplay between potentially opposing factors such as microbial resource use efficiency, resource acquisition and resource availability, itself depending on SOM and plant input quantity and quality (e.g. root exudates). Moreover, microbial biomass depends on the whole soil trophic network (Calderón-Sanou et al. 2021). Furthermore, as found in alpine ecosystems in Cordillera Darwin, Tierra del Fuego (Chile) (Thébault et al. 2014), we suggest that competition between soil microbes and plants for nutrient resources in SOM and TDN-rich soils may contribute to the higher mass-specific microbial investment in resource acquisition in these soils. This result stresses the importance to consider the resource acquisition strategy of soil microbes and not only their biomass to understand their response to environmental gradients and their effect on nutrient recycling in soil (Piton et al. 2020a). Recent literature has proposed that microbes with an A strategy could be favored either when resources are scarce (Malik et al. 2019, 2020; Piton et al. 2020b), or when resources are abundant (Wood et al. 2018). Since microbes rely on two main carbon sources, not only soil organic matter but also those exuded by plants in the rhizosphere, it is necessary to explore the relationships between the traits of the microbial and plant communities. We found that microbes increased their investment in extracellular enzymatic activities in plant communities having higher leaf C/N ratios (Figure 1C), which corresponds to conservative plant resource use strategy. Since exploitative plants have been shown to produce more root exudates (Williams et al. 2022), an extra labile carbon source would be available to microbes in exploitative plant communities, favoring microbes with a high yield (Y) strategy characterized by low enzyme production, according to Malik et al. (2020). This might also be reinforced by organic inputs in the most productive intensive grasslands. In contrast, in conservative plant communities, microbes would lack labile carbon and nutrients sources and rely more on the biodegradation of polymerized soil organic matter for resource acquisition. This implies that A strategists characterized by the production of extracellular enzymes would be favored, which ultimately releases nitrogen from soil organic matter (Figure 1C).

**Table 2.**
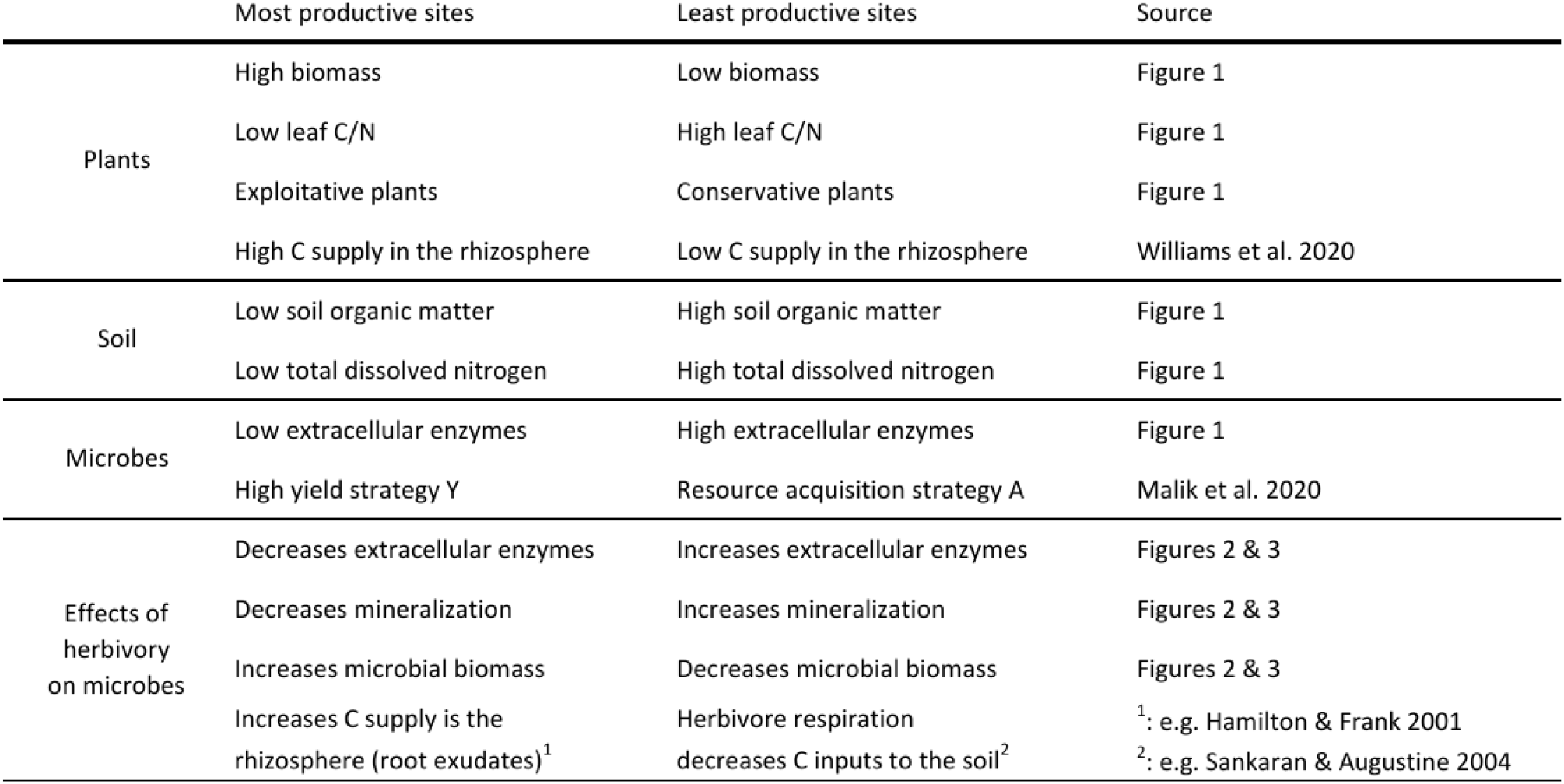
Schematic representation of the main results, and their articulation with previous works.

In summary, the pattern of covariations between ecosystem properties suggests a distinction between two kinds of grasslands. Although both microbial strategies are likely to coexist in each soil sample, both grassland kinds favor either strategy. More specifically, on the one hand the most productive sites (>400 g dw/m^2^) are characterized by exploitative plants, by soils having relatively low SOM (<10%) and TDN (<40 μgN/g) contents, and by Y-strategist microbes producing fewer extracellular enzymes, possibly because of higher C supply in the rhizosphere (Table 2, left column). On the other hand, in the least productive sites dominated by conservative plants, the SOM and TDN contents are higher and the microbial community is characterized by an A strategy since microbes need to produce extracellular enzymes to get access to SOM resources (Table 2, right column).

### Herbivory effects on enzymatic activities

The present section explores if the effects of herbivory on microbial communities depend on the two grassland kinds described above (Table 2). We found a covariation between SOM and the responses of soil enzymatic activities and PNM to herbivory (Figure 2). Herbivory had a negative effect on soil enzymatic activities and PNM when SOM was below 10%, and a positive effect when SOM was above 15% (Figure 3), which means that herbivory can decelerate or accelerate nutrient cycling depending on soil conditions. This occurs in relation to microbial strategies; in this section we interpret the variations of soil enzymatic activities following herbivory as responses of the microbial community, favoring either A or Y strategy. In soils rich in SOM already characterized by high mass-specific extracellular enzyme activities, microbes invested even more in extracellular enzymes in response to herbivory indicating that A strategists were favored under these conditions. Herbivory provides fresh organic matter (FOM such as green fall, feces, cadavers) to the soil, which contains nutrients that are more easily available than SOM but which nevertheless needs to be depolymerized before assimilation (e.g. chitin). Since in rich-SOM-sites, A strategists possess the enzymatic traits that give them access to this resource, this would explain why herbivory promoted the investment of A strategists in extracellular enzymes. Consequently, the enhancement of extracellular enzymatic activities might have cascaded on PNM, as expected according to the key limiting role of organic matter depolymerization before microbial assimilation and subsequent N mineralization (Schimel and Bennett 2004). In contrast, in low SOM soils, Y strategists decreased even more their investment in extracellular enzymes. A possible explanation is that litter inputs are not the only consequences of herbivory on ecosystem functioning (Hunter et al. 2012). In particular, herbivory stimulates root exudation (Holland 1995; Holland et al. 1996), which is known to increase microbial activity in the rhizosphere and subsequently to favor compensatory plant growth (Hamilton and Frank 2001; Hamilton et al. 2008). Williams et al. (2022) found that root exudation is more intense in exploitative than in conservative plant communities, but it is not known whether the effect of herbivory on root exudation depends on plant functional groups. If the stimulation of root exudation by herbivory is more intense for exploitative than for conservative plants, this would explain the finding that herbivory decreased the production of extracellular enzymes in sites having low SOM, since they were dominated by exploitative plants. However, although the effect of herbivory on soil enzymatic activities and PNM depended on SOM content (Figure 3), it did not directly depend on the mean leaf C/N ratio (details not shown, p>0.05). Root exudation following herbivory has been found to enhance PNM for the benefit of plants (Hamilton and Frank 2001), while our results suggest the opposite. However, inputs of available C may also result in N immobilization in the microbial biomass (Lovett and Ruesink 1995), which is this time in line with our results. This calls for both qualitative and quantitative characterization of root exudation in future works to further elucidate how soil and plant properties jointly control ecosystem response to herbivory. Herbivory may also decelerate nitrogen mineralization through the induction of recalcitrant plant secondary compounds (Schultz and Baldwin 1982; Rhoades 1985; Agrawal 1999), which subsequently slows down litter decomposition (Findlay et al. 1996; Hattenschwiler and Vitousek 2000). However, we did not find any effect of herbivory on litter decomposition in a parallel study (Ibanez et al. 2021).

Turning to microbial biomass, herbivory had a positive effect in sites having low TDN, and a negative effect when TDN was high (Figure 3E), but contrary to the enzymatic activities the effect of herbivory on microbial biomass did not depend on the SOM content (Figure 2A). In previous studies, herbivory increased microbial biomass through an input of labile C contained in the feces (Lovett and Ruesink 1995; Van Der Wal et al. 2004). Yet, herbivory was also found to decrease microbial biomass when plants’ C inputs to the soil are instead reduced due to herbivores respiration (Sankaran and Augustine 2004). Since TDN was positively linked with mean leaf C/N ratio (Figure 1B, Robson et al. 2007; Legay et al. 2013), itself being negatively correlated to root exudates (Williams et al. 2022), we hypothesize that in low TDN sites, herbivory triggered an input of labile C into the rhizosphere, thereby stimulating the growth of microbial biomass, whereas in sites with high TDN, herbivory reduced the plants C inputs, with a negative impact on microbial biomass.

### No effect of herbivore diet

The proportion of forbs in the diet of *M. alpina* was the highest, followed by the mixed feeder *S. scalaris* and then by the grass feeder *P. parallelus* (Sup. Fig. 2A), in line with previous studies (Ibanez et al. 2013). Since the grasshoppers’ diet did not depend on the grasslands they were introduced in, it was possible to test if the effects of herbivory on soil enzymatic activities depended on their diet. The diet model predicts that herbivores feeding on high quality plants should favor the accumulation of poor-quality litter and thereby slow down nutrient cycling, and that herbivores feeding on low quality plants should favor the accumulation of high-quality litter, with an accelerating effect on nutrient cycling (Belovsky and Slade 2000).

In the present study, high quality plants characterized by low leaf C/N ratio (Sup. Fig. 4) were consumed preferentially by the forb feeder *M. alpina* (Sup. Fig. 2A), which decreased their relative abundance by about 7% (Sup. Fig. 5). In warm and forb-dominated prairies, litter decomposition during winter has been found to be faster for forbs (28% of mass loss) than for grasses (17%), although in extensive and intensive grasslands winter decomposition was similar for both plant functional groups, due to the intermittent presence of snow cover (Ibanez et al. 2021). In any case, this suggests that the year-round decomposition rate is higher for forbs than for grasses, as generally described (Tilman 1988). Given that *M. alpina* modifies the balance between forbs and grasses, this should affect the overall decomposition rate, with potential impacts on soil microbial communities. However, *M. alpina* did not have any contrasted effect on soil microbial characteristics, in comparison to the other grasshopper species. Perhaps this would have required a heavier impact on the relative abundance of forbs, rather than only 7%.

Lower quality plants (high leaf C/N ratio) were consumed preferentially by the mixed feeder *S. scalaris* and almost exclusively by the grass feeder *P. parallelus* (Sup. Fig. 2A), but without impact on the relative abundance of forbs and grasses (Sup. Fig. 5). Also, grass feeders did not have any distinguishable effect on soil microbes, relatively to the forb feeding *M. alpina*. Previous short-term experiments using similar insect loads have reported that the effects of grasshoppers on soil processes depended on their diet type (e.g. Schmitz 2008; Belovsky and Slade 2018). Our findings question the generality of these results, since we found that the effect of herbivory on enzymatic activities did not depend on grasshopper species identity.

## Conclusion

Previous works showed that herbivory has contrasted impacts on ecosystem functioning in general (Brown and Gange 1992; Bardgett 2005; Schmitz 2008), and on soil microbes in particular (Denton et al. 1998; Stark and Grellmann 2002), in function of the environmental conditions and of the diet of herbivores. The motivation of this work was to observe such contrasted impacts in a single experiment across different types of grasslands with contrasting plant productivity, using herbivores having different diets, in order to better understand the pathways which create this apparently idiosyncratic pattern. We did not find any effect of herbivore diet on soil microbial characteristics, contrary to our expectation. In contrast, the effects of herbivory on soil microbes depended on several properties of the 12 study sites. Table 2 summarizes our main findings and can provide some lines of interpretation, although its dichotomic schematization does not fully represent the multidimensionality of the results (Figure 1–3). On the one hand, the most productive sites were characterized by a higher biomass of exploitative plant species which depleted N resources in the soil, and by yield-strategists microbes having a smaller investment in extracellular enzymes. Since exploitative plant species tend to produce more root exudates (Williams et al. 2022), we postulate that herbivory increased C supply in the rhizosphere (e.g. Holland et al. 1996; Hamilton and Frank 2001), which would explain the observed increase in microbial biomass and the lesser production of extracellular enzymes. On the other hand, the least productive sites were characterized by a lower biomass of conservative plants, which led to high soil C content, and by acquisition-strategists microbes having a larger investment in extracellular enzymes. We hypothesize that, in those sites, the consumption of plants results in lower soil C inputs due to herbivore respiration (Sankaran and Augustine 2004), which would explain why herbivory eventually decreased microbial biomass and increased the investment in extracellular enzymes in these less productive sites. Although the framework presented in Table 2 has some similarities with the productivity model (Bardgett and Wardle 2010), our findings point towards an acceleration of N cycling in less productive sites and a deceleration in more productive sites, in opposition with the productivity model. However, both frameworks do not consider the same time scales. The current experiment was conducted over two years and focuses on physiological time scales (root exudation, microbial enzyme production), while the productivity model encompasses plant community dynamics, which occur on longer time scales. In any case, we are convinced that none of these frameworks fully grasp the complex relationships between plants, soil, microbes and the effects of herbivory, which are most likely multidimensional. At the very least, these frameworks have some heuristic value and can be used for the design of future experiments.

## Acknowledgements

This work was funded by the ECO-SERVE project through the 2013–2014 BiodivERsA/FACCE-JPI joint call for research proposals, with the national funders ANR, NWO, FCT (BiodivERsA/001/2014), MINECO, FORMAS and SNF. This work was also funded by the Alpine Ecology Lab. We thank the municipality of Autrans-Méaudre, the local farmers and the Refuge de Gève for their authorization to access the study sites, Jonathan Crison for his hospitality. We are grateful to Jean-Noёl Avrillier, Annie Millery, Hugo Girard, Matteo Tolosano, Louise Maris, Pablo Raguet, Océane Guillot, Jessica Barbe & Alison Dillien for their assistance during field and lab work.

## Supplementary Figures & Tables

**Sup. Fig. 1.**
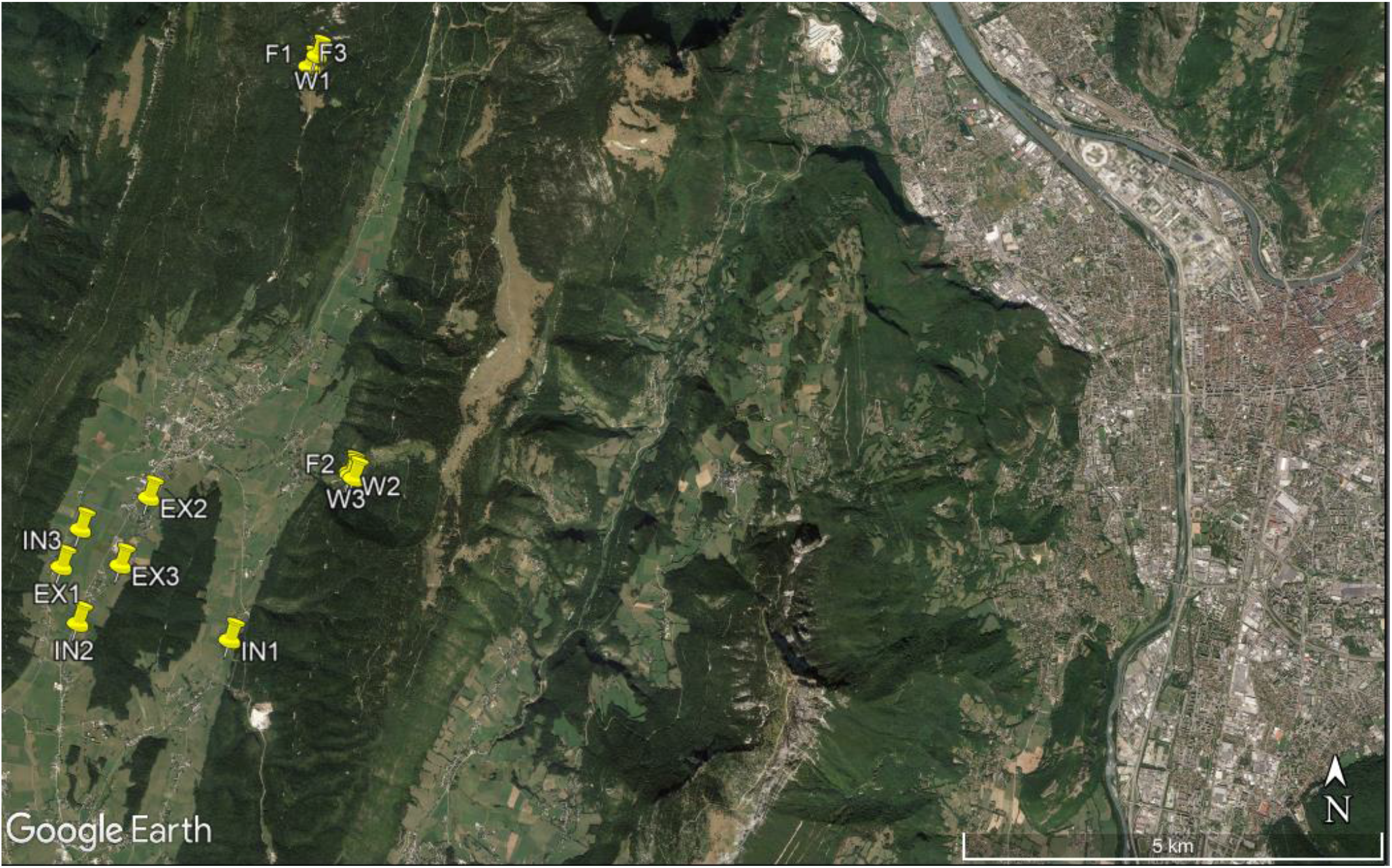
Geographical distribution of the 12 study sites (F = Forbs communities, W = Warm grasslands, IN = Intensive grasslands, EX = Extensive grasslands). F1, W1 and F3 are close to each other, as well as F2, W2 and W3. Grenoble corresponds to the large city at the east.

**Sup. Fig. 2.**
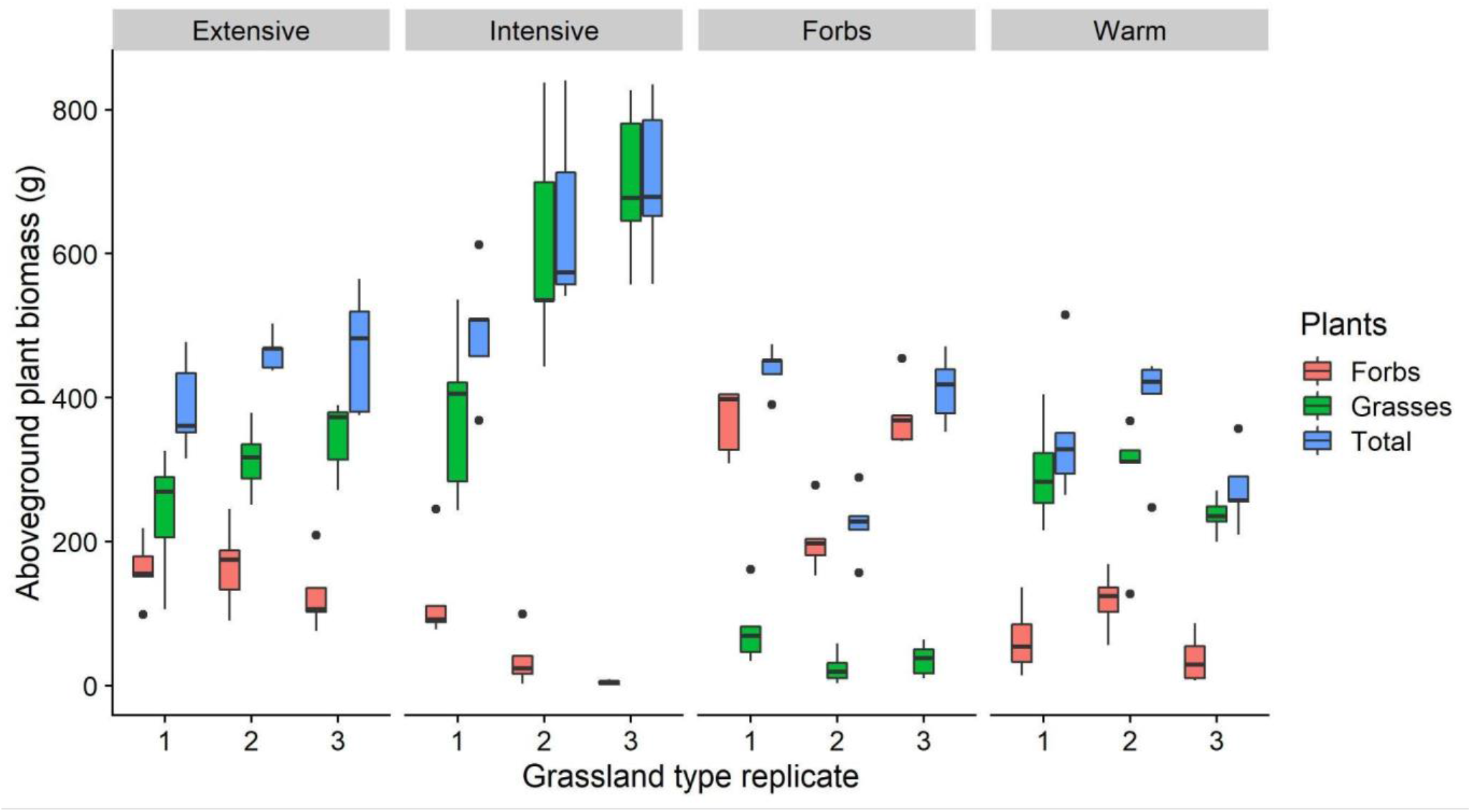
Aboveground plant biomass for grasses and forbs, in function of grassland type. There are 12 sites, 4 grassland types and 3 replicates per type. Each site is composed of 5 1m^2^ cage.

**Sup. Fig. 3.**
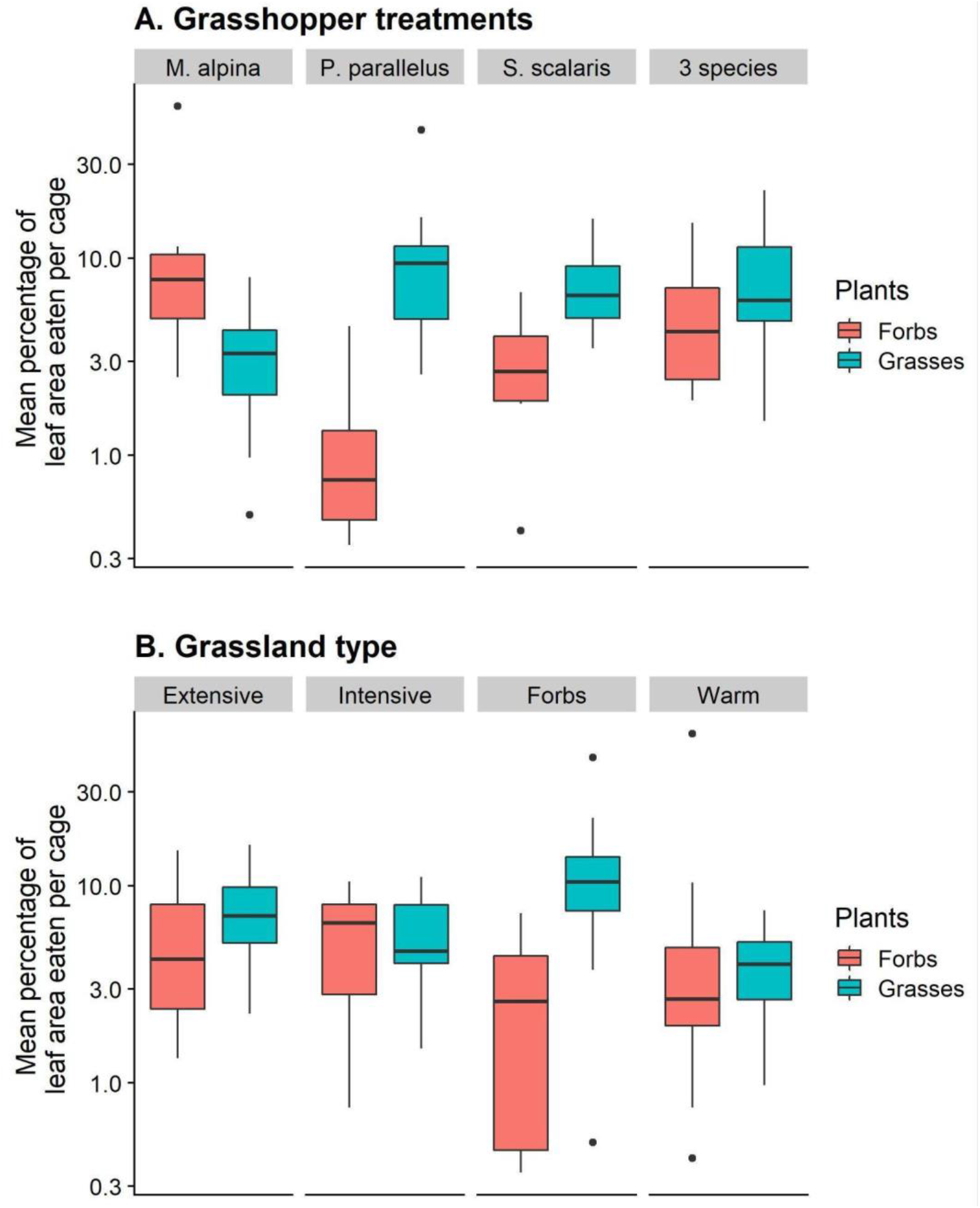
Mean percentage of leaf area eaten for each plant functional group, in function of (A) grasshopper treatment, and (B) grassland type. The percentage of leaf area eaten was estimated from observations (mid July 2017) of the dominant plant species in each plot.

**Sup. Fig. 4.**
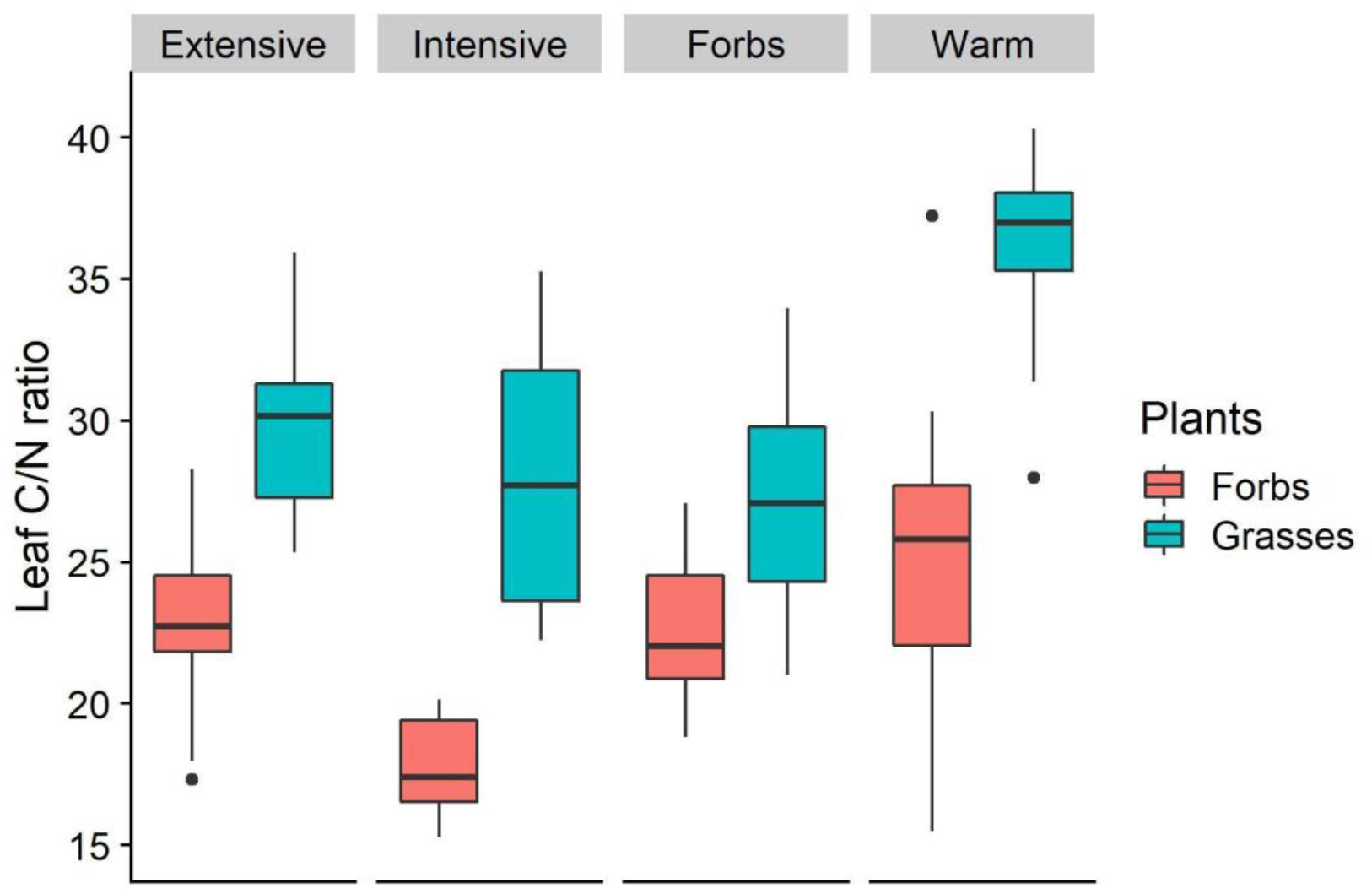
Leaf C/N ratio of forbs and grasses in the four grassland types.

**Sup. Fig. 5.**
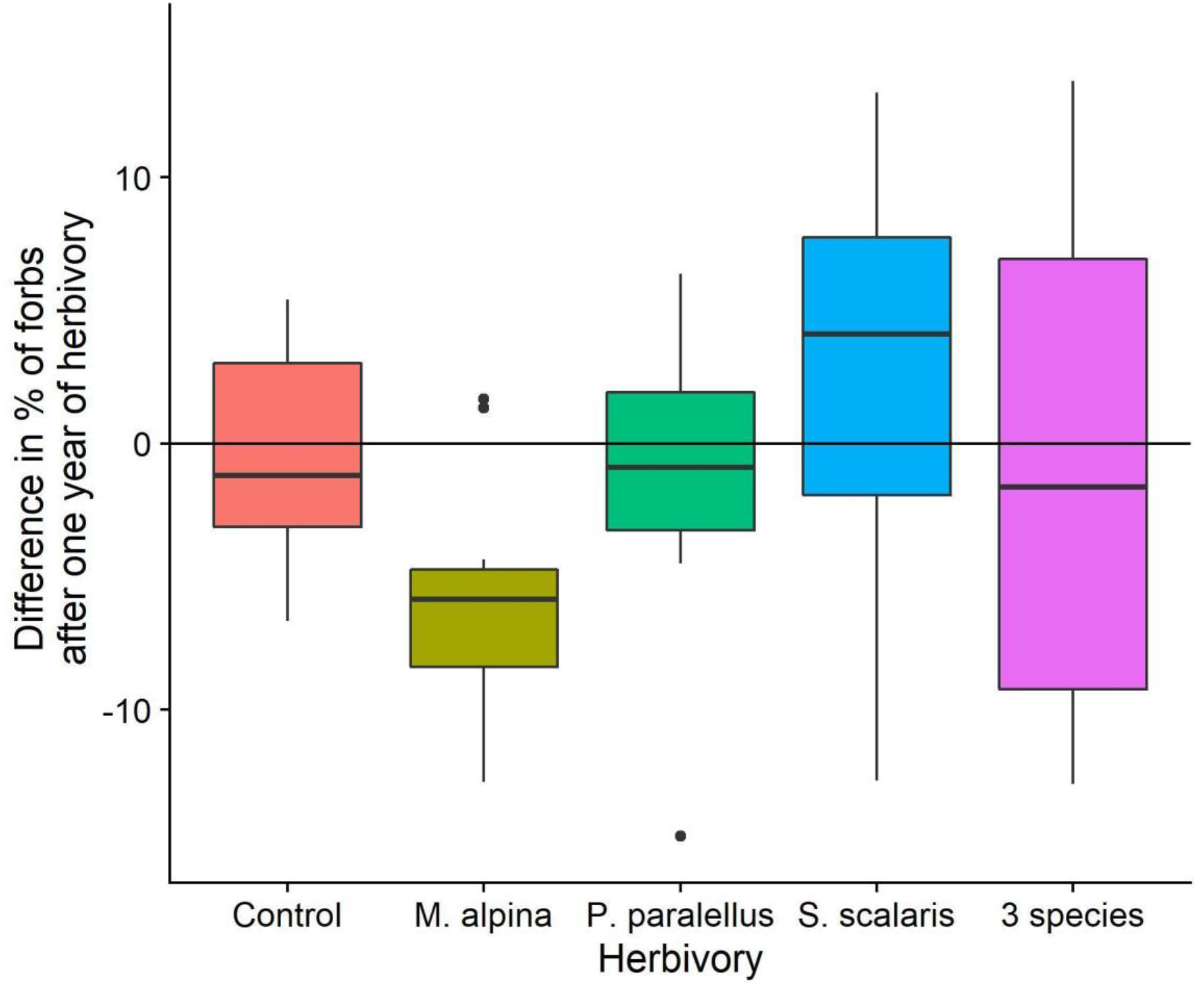
Variation of the percentage of forbs estimated by the point quadrat method after one year of herbivory, in function of the herbivory treatment.

**Sup. Fig. 6.**
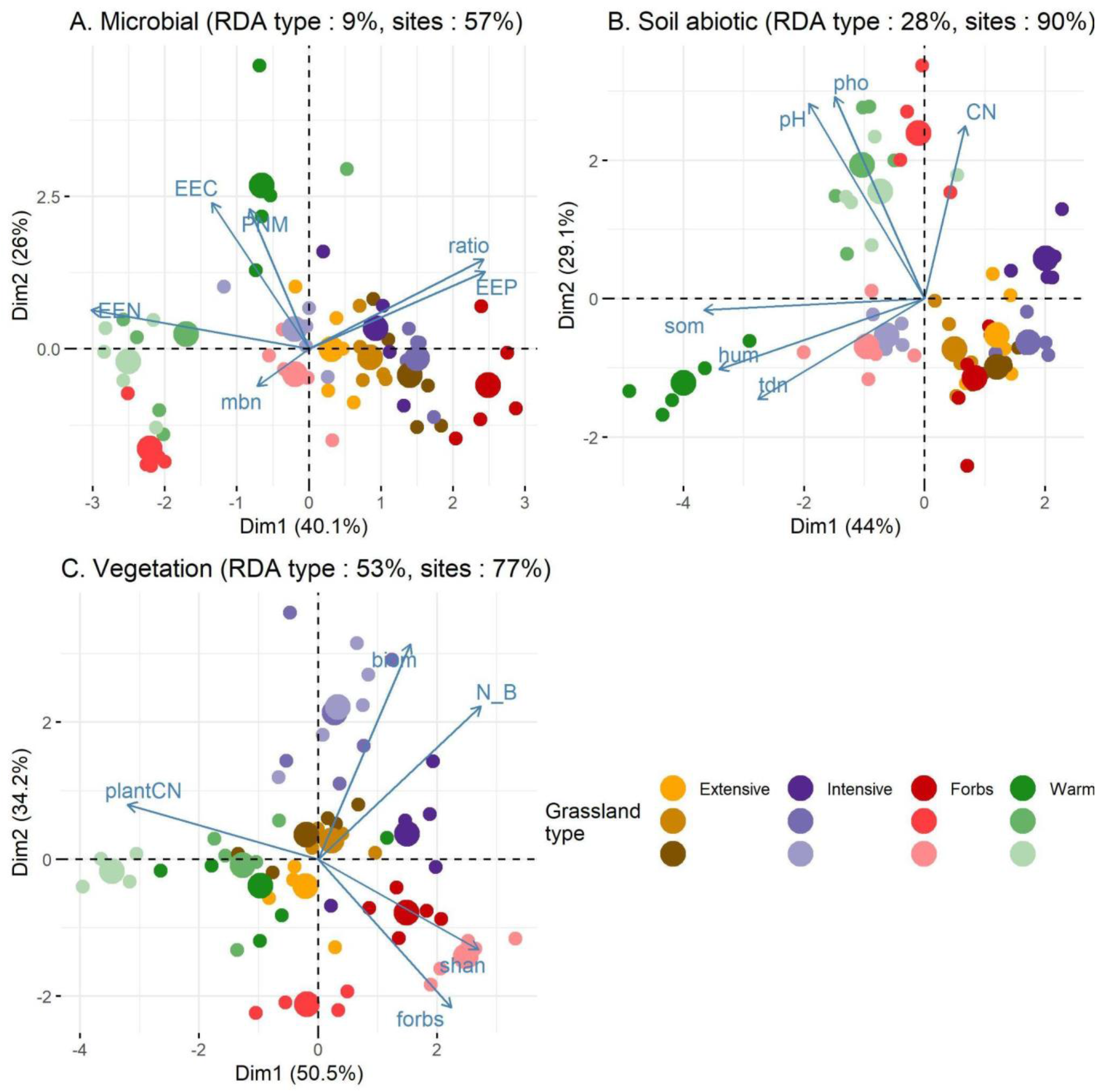
Principal component analysis of A. microbial, B. soil abiotic and C. vegetation characteristics. The percentage of variation explained by either the 4 grassland types or the 12 sites are given (redundancy analysis).

**Sup. Tab. 1.**
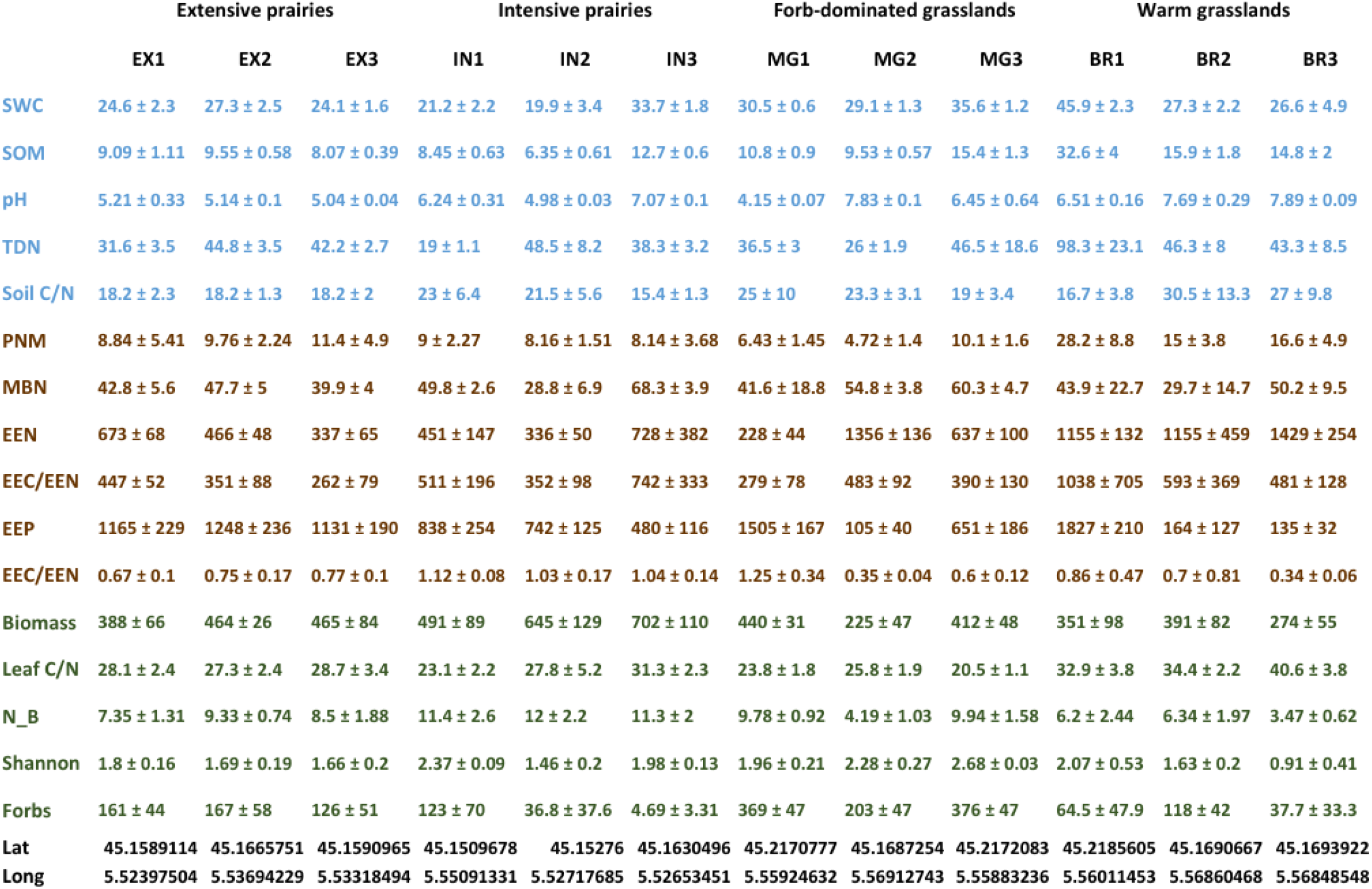
Mean±sd of each of the 12 sites (with 5 plots per site). <u>SWC</u>: Soil water content (%), <u>SOM</u>: Soil organic matter (%), <u>TDN</u>: Total dissolved nitrogen (μg/g), <u>PNM</u>: Potential nitrogen mineralization (μgN/g dry soil/day), <u>MBN</u>: Microbial biomass nitrogen (μg/g), <u>EEN</u>: Nitrogen-related enzymes (nmol activity/g dry soil/h), <u>EEC</u>: Carbon-related enzymes, <u>EEP</u>: Phosphorous-related enzymes, <u>Biomass</u>: Plant biomass (g/m^2^), <u>N B</u>: Nitrogen in aboveground plant biomass (g/m^2^), <u>Forbs</u>: Forb biomass (g/m^2^). Geographical coordinates are also given.

## Data accessibility

Data used for the analysis: https://doi.org/10.5061/dryad.5tb2rbp6j

R script used for the analysis: https://doi.org/10.5281/zenodo.6831330

